# Multi-scale disruption of sleep-related cortical activity following ischemic lesions

**DOI:** 10.64898/2025.12.19.695488

**Authors:** Vinícius Rosa Cota, Simone Del Corso, Federico Barban, Marta Carè, Michela Chiappalone

**Affiliations:** Department of Informatics, Bioengineering, Robotics and Systems Engineering (DIBRIS), Università degli Studi di Genova, Genova, Italy; Department of Electronic Engineering, Maynooth University, Maynooth, Ireland; Rehab Technologies Lab, Istituto Italiano di Tecnologia, Genova, Italy; IRCCS Ospedale Policlinico San Martino, Genova, Italy

## Abstract

Sleep plays a critical role in neuroplasticity related processes of learning and memory consolidation. Moreover, its importance in recovery after stroke is known. Yet, how ischemic lesions affect the sleep-wake cycle (SWC) neural dynamics remains poorly explored, with most studies not addressing the multi-scale extent of this pathology effects in brain function. Here, we performed chronic wideband electrophysiological recordings in behaving rats to investigate ischemic stroke impact in the primary motor cortex on SWC organization in connected, non-lesioned, areas of the sensorimotor loop. We combined behavioral assessment with multi-scale analyses spanning global SWC architecture, network level events of privileged functional reshaping during sleep, and single-unit activity. Lesioned animals, exhibiting sharp motor impairments, displayed clear alterations of SWC macro-architecture, including reduced latency to slow-wave sleep and correlations between spectral features and motor asymmetry. At meso-scale, ischemia disrupted the physiological coordination between slow oscillations and spindles, and the latter showed shift in spectral power distribution during intermediate sleep. In contrast, units firing and coupling with spindles were largely preserved. These findings reveal that ischemia perturbs the fine coordination of sleep-related processes across scales stage-specifically. The study lays solid evidence to be possibly leveraged for innovative protocols of neurorehabilitation targeting privileged windows of neuroplasticity.

## 1. Introduction

Motor impairment following stroke is the leading cause of late-onset disability and a major cause of death worldwide (Campbell et al. 2019). First choice treatment for stroke survivors is usually carried out as physical therapy aimed at motor rehabilitation (Coleman et al. 2017), while robotic-aid and pharmacological options are mostly adjunctive. A promising alternative treatment relies on neurostimulation, aimed at promoting brain plasticity (neuroplasticity) and improving functions (Bao et al., 2020; Saway et al., 2024; Carè et al., 2024). Yet, a significant proportion of patients still fall short of re-acquiring adequate level of motor function for daily-life activities.

It is now widely recognized that sleep plays a major role in the mechanisms underlying neuroplasticity (Peigneux et al. 2001; Walker and Stickgold 2006; Lanza et al. 2022). Consolidation of recently acquired memory traces and motor skills are largely dependent on neurobiological mechanisms provided in a complementary way by the different brain states (or stages) across the sleep-wake cycle (SWC) (Dang-Vu et al. 2006; Brodt et al. 2023; Ribeiro 2012; Vahdat et al. 2017). Notably, there is now mounting evidence that precise temporal coordination between cortico-cortical slow oscillations, thalamocortical spindles, and hippocampal ripples supports the reverberation of memory traces-related activity (Ribeiro et al. 2004; Born and Wilhelm 2012; Girardeau et al. 2009; Girardeau and Lopes-Dos-Santos 2021; Diekelmann and Born 2010), which then guides neurochemical cascades for due plastic changes (Ribeiro et al. 1999). In this framework, spindles seem to be of major importance for motor learning and recovery, as they mediate the synchronization across networks involving basal ganglia and the cortex (Boutin et al. 2018) or across cortical regions (Darevsky et al. 2024). This takes place mostly during stage 2 of non-REM (Rapid Eye Movement) sleep in humans (Boutin et al. 2024) or intermediate sleep (IS; between Slow Wave Sleep - SWS - and REM) in rodents (Blanco et al. 2015).

It is clear that understanding whether and how ischemic lesions reshape sleep-related cortical dynamics across global, network, and unitary activity scales, in a stage specific fashion, can help in designing novel rehabilitation therapies aimed at favoring post stroke recovery (Duss et al. 2017). Electrographically, stroke is associated with significant changes, which include widespread arrhythmic delta activity, reduction of ipsilateral alpha rhythm, and emergence of epileptiform discharges in humans (Niedermeyer 1982). At the network level, experimentally induced stroke leads to a higher occurrence of isolated delta waves and disrupts the temporal coupling between Slow Oscillations (SOs) and spindles in the perilesional cortex, as recently reported in (Kim et al. 2022). Pharmacological reduction of GABAergic function in that study was observed to suppress such a dysfunctional activity which was correlated to functional recovery as well.

Despite the recognized bidirectional relationship between sleep and stroke (Khot & Morgenstern, 2019), the specific impact of cortical ischemic lesions on the architecture and dynamics of the SWC remains poorly understood, particularly in terms of how lesion-induced alterations manifest across the hierarchical organization of brain activity. Here, we address this scientific gap by performing a comprehensive, multi-scale electrophysiological investigation in chronically implanted animal models. We focused our attention on the sensorimotor loop, a possible target of neurotechnological approaches for motor rehabilitation (Guggenmos et al. 2013). Specifically, we induced a focal lesion in the caudal forelimb area – CFA (i.e. the equivalent of the motor cortex in humans) and evaluated the electrophysiological activity of both the pre-motor area (rostral forelimb area; RFA) and the sensorial cortex (primary somatosensory cortex; S1), two brain regions which are not directly impacted by the lesion itself but are directly connected to the damaged area.

We assessed lesion-induced changes in global oscillatory patterns and SWC states (macro-architecture), in the expression of sleep-related network events such as spindles and slow waves (meso-architecture), and in the underlying neuronal firing dynamics (micro-architecture) at the cortical level, as well as their cross-scale interactions over time (micro-to-meso and meso-to-macro coordination). Given the potentially preponderant role of spindles occurring during IS, we assessed the impact of lesion also in this perspective. We found that the ischemic lesion induces widespread multi-scale disruption of neural activity across the SWC, from the macro down to the micro-architecture.

Our findings highlight the need for a multiscale electrophysiological framework to reveal how surviving circuits reorganize after ischemic injury, providing a foundation for neurotechnologies that aim to boost plasticity and promote functional recovery post damage.

## 2. Material and Methods

### 2.1. Animals and groups

Thirteen (N = 13) Long Evans male rats (*Rattus norvegicus*), weighting 250–300 g, obtained from Charles River (Calco, LC, Italy), were used in this study. They were housed at the vivarium of IIT’s Animal Facility for the entire duration of the experiment, being kept in a 12-hour light–dark cycle (lights on at 7 am and off at 7 pm), at an average temperature of 23°C ± 2°C, with food and water ad libitum. Animals were divided into two groups, controls (CTRL; N = 7) and lesion (LESION; N = 6). All animals underwent surgical procedures for the implantation of electrodes in cortical regions of the sensorimotor loop (i.e., RFA and S1; Figure 1A-B). Animals in the LESION group were also subjected to the induction of an ischemic lesion (*cf.* section 2.2 for details). After a 7-day recovery period, all animals underwent six hours-long electrophysiological recordings followed by behavioral testing for assessment of motor function. All experiments were previously approved by the Italian Ministry of Health (license 513/2022) and are in accordance with international guidelines for the care of animals in research.

**Fig. 1.**
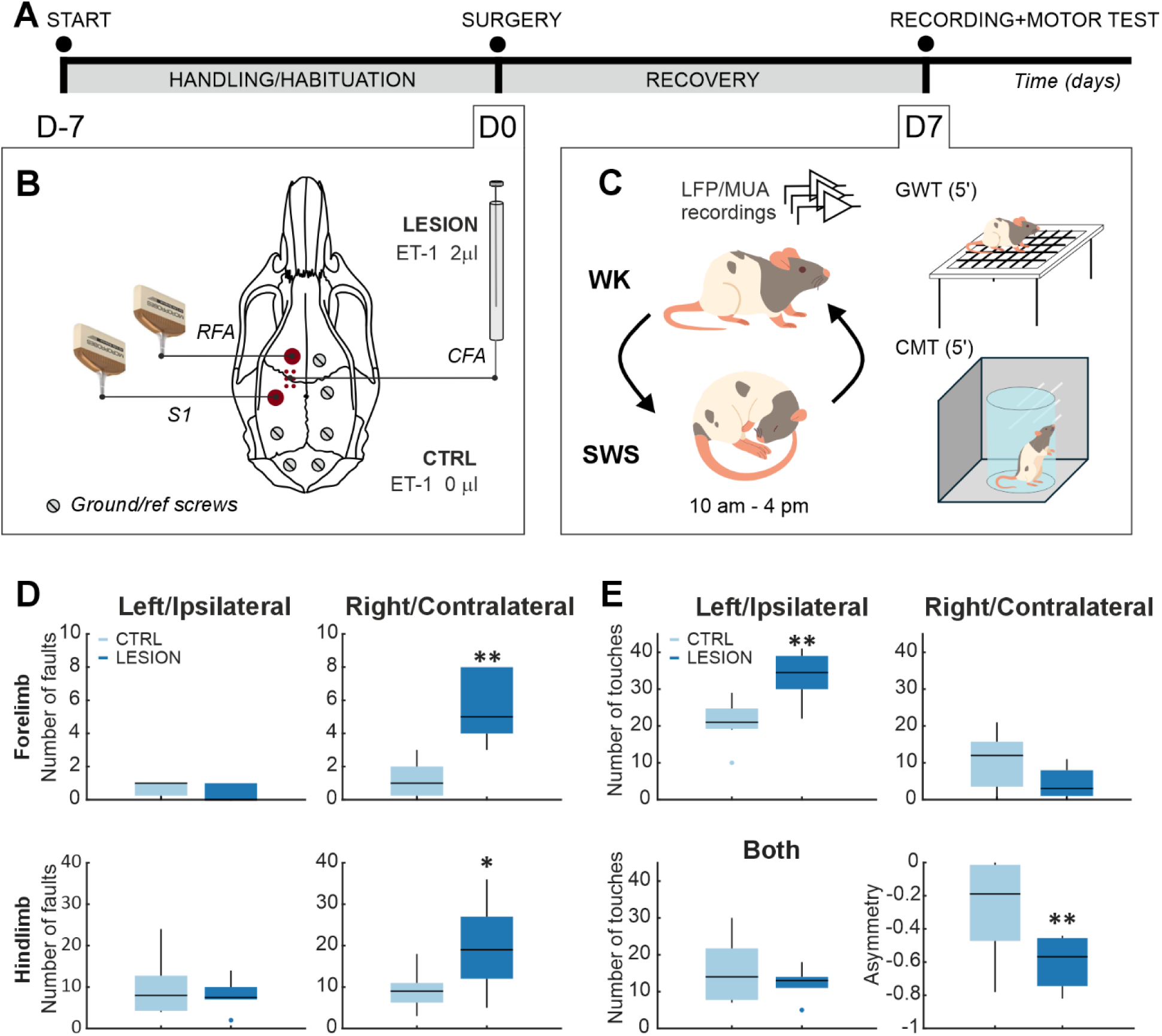
Experimental protocol and behavioral assessment. (**A**) Experimental timeline starts seven days before surgical procedure (D-7) with the handling and habituation of the animals. After surgery at day ‘zero’ (D0) for the implantation of recording electrodes and induction of lesion (LESION group only, differently from the CTRL animals that were not lesioned), animals were allowed to recover for 7 days and were then submitted to electrophysiological recordings and testing of the motor function (D7). (**B**) Surgical procedures were performed on D0 for the implantation of two 32 channel microwire arrays in the Rostral Forelimb Area (RFA) and the primary somatosensory area (S1) and for the induction of an ischemic lesion (LESION group only) by means of microinjection of ET-1 in the Caudal Forelimb Area (CFA). Six stainless steel surgical microscrews were inserted in the animal cranium for ground and referencing, as well as for anchoring the whole implanted assembly. (**C**) On D7, animals were first submitted to electrophysiological recordings of Local Field Potential (LFP) and Multi Unit Activity (MUA) for 6 hours (from 10 am to 4 pm), during which they were able to freely move around (Wakefulness state, WK) or sleep (Slow Wave Sleep state, SWS). Later, animals were submitted to the Grid Walking Test (GWT) and the Cylinder Mirror Test (CMT) in a 5-minute session each. (**D**) Number of full slips across openings in the GWT with each of the four limbs, fore and hind, left (ipsilateral) and right (contralateral). Animals in the LESION group displayed a significant higher number of slips with both right forelimb (top right; **p = 0.002, Mann Whitney U test) and right hindlimb (bottom right; *p = 0.037, Mann Whitney U test). There were no significant differences between groups for both fore and hindlimbs of the ipsilateral side. (**E**) Results of the CMT showing the number of touches in the cylinders walls with the left (ipsilateral), right (contralateral), or both paws simultaneously, as well as the computed asymmetry. Animals in the LESION groups displayed a significantly increased number of touches with the left paw (top left; **p = 0.006, Mann Whitney U test) as well as a significantly lower asymmetry index (bottom right; *p = 0.037, Mann Whitney U test), correlating to a more intense usage of the left paw. In all panels of this figure, CTRL animals are depicted in unsaturated blue and LESION in saturated blue colors.

### 2.2. Surgical procedure for electrode implant and induction of ischemic stroke

Prior to any surgical procedure, initial sedation was induced with the administration of gaseous isoflurane (5% at 1 lpm) using a dedicated inhalation chamber for the animal. This was followed by the administration of ketamine (intraperitoneal, 80-100 mg/kg) and xylazine (intramuscular, 5 mg/kg). After trichotomy of the head, protection of the eyes with ophthalmic ointment, and appropriate aseptic measurements with Betadine and 70% alcohol, animals were positioned in the stereotaxic apparatus. Body temperature was monitored by a rectal probe and controlled by a heat pad using a physiological monitoring system, Physio Suite® (Kent Scientific, Torrington, U.S.A). The sedation status of the animal was continuously monitored by the experimenters, checking for animal twitches through a simple pinch test and, when necessary, maintenance boluses of ketamine (intramuscular, 10-20 mg/kg) were administered. The skull surface was exposed with a rostro-caudal incision using a scalpel, followed by dissection of the periosteum, and cleaning of the bone with hydrogen peroxide. In the sequence, the cisterna magna was exposed, and a laminectomy was performed to control for brain edema, by allowing excess cerebrospinal fluid drainage. Craniectomy was then performed to access the primary somatosensory cortex (S1: -1.25 mm AP and +4.25 mm ML, referenced from the Bregma) and the rostral forelimb area (RFA, a pre-motor area: +3.5 mm AP and +2.5 mm ML, referenced from the Bregma) using a drill with a burr bit (Figure 1B). Additional six holes were predisposed with a small drill bit into the parietal and intraparietal bones to later insertion or surgical microscrews that served both as reference and grounding, as well as to secure the implant on the animal for the chronic recordings. Dura-mater was carefully removed using a bent hypodermic needle to avoid excessive brain tissue dimpling and electrode bending and damage. Electrodes (two 32 channel microwire arrays, one for each area; Microprobes, USA) were slowly (100 μm/min) inserted into the cortex to a depth of 1.4 mm referenced from the cortical surface. Implantation sites were sealed using low toxicity adhesives for live tissues (Kwik-cast; WPI, Sarasota, USA). The whole assembly, including both arrays, was secured onto the animal’s head with dental auto polymerizing acrylic cement anchored by the bone microscrews. Finally, the skin opening was sutured, animals received analgesic and anti-inflammatory treatment (Diclofenac sodium: Voltaren, 5 mg/kg s.c.), as well as antibiotics (Enrofloxacin: Baytril, 10 mg/kg), were cleaned, and remained under observation in a heated cage until they woke up. Post-surgery treatment with painkillers, antiinffamatory, and antibiotics was repeated in the following two days.

Animals in the LESION group were also submitted to the ischemic stroke model by microinjection of ET-1, a potent vasoconstrictor peptide (Frost et al. 2006)(Figure 1B). During the surgical procedure, before the insertion of electrode arrays, six small holes were drilled in the skull centered around the caudal forelimb area (CFA), equivalent to human primary motor cortex (combining coordinates at AP: +1.5 mm, +0.5 mm, and -0.5 mm; ML: +3.5 mm and +2.5 mm, referenced from Bregma). ET-1 was then loaded into a Hamilton syringe which was lowered 1.2 mm into brain tissue. A volume of 0.33 μl per hole (thus a total of 2 μl) was injected, being divided into three steps of 0.11 μl each, gradually administered along 1 minute and interspaced by also 1 minute. This guaranteed a seamless absorption of the vasoconstrictor agent volume without damage by tissue displacement.

### 2.3. Behavioral testing of motor function

To evaluate the impact of the ischemic lesion on the motor function of animals, two well-established tests were used. Imposed and spontaneous movement functions were respectively assessed by, respectively, the grid walking test (GWT) and the cylinder-mirror test (CMT), (Figure 1C). In the GWT (Barth et al. 1990), animals are put in an elevated square grid of circa 50 x 50 cm with bars interspaced circa 2.5 cm in both directions and were allowed to freely explore it for 5 minutes. The number of faults, consisting of full slips across the openings, were then counted for each limb. It is expected that animals suffering from motor dysfunction will display increased numbers of faults in the contralateral limb in respect to the lesion side (in our case, right limbs). In the CMT, animals were inserted inside a plexiglass cylinder of 30 cm heigh and circa 20 cm in diameter positioned at the side of two perpendicular mirrors and were left to explore it also for five minutes. The number of touches in the cylinder wall with the left, right, and both forepaws simultaneously, was counted. It is expected that lesion animals will display a greater tendency to use the ipsilateral non-impaired paw. Several different metrics can be computed for an objective assessment of this effect (Schallert 2006; Schallert et al. 2000). Among them, the asymmetry index (*A*) computed from the percent impaired forelimb touches subtracted from the percent nonimpaired forelimb was chosen here. It is obtained by the following (Eq. 1):

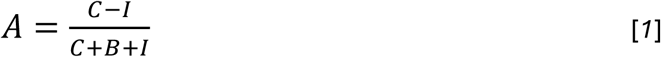

Where *C* is the number of wall touches using the contralateral (i.e., impaired) paw, *I* by the ipsilateral (i.e., nonimpaired) paw, and *B* by both paws. While healthy animals display asymmetry indexes close to zero, lesion ones tend to induce an asymmetry of limb usage towards the ipsilateral paw.

In both GWT and CMT, animals were naïve to the apparatuses. In order to lower stress levels during experimental sessions, all animals were handled by the experimenter for 15 minutes for five days, twice a day, before surgery (Figure 1A). Behavioral tests were always performed after recording sessions, starting at around 4.30 pm and were filmed on digital video.

### 2.4. Electrophysiological recordings and signal pre-processing

In the day of the recording session (i.e., D7), animals were lightly anesthetized with gaseous isoflurane (5% at 1 lpm) for cleaning of electrode array contacts and for the placement of recording headstages and cables. Animals were then placed in the arena and cables connected to the electrophysiological setup. The system was composed of a controller (Intan Technologies LCC, Los Angeles, USA) for host PC communication and acquisition control, connected via SPI to headstages (equipped with RHS 2000 series chips; Intan Technologies LCC, Los Angeles, USA) that performed amplification, filtering, and digitalization. Biopotentials were initially filtered by a light analog 1-pole high pass filter at 0.1 Hz cutoff frequency (DC decoupling) and then digitally at 1 Hz (as per usual configuration of the Intan system; see for instance (Saleem et al. 2010)). An anti-aliasing 8^th^ order low pass filter with 7.5 kHz cutoff frequency was also applied before amplification (192 V/V) and sampling at 25 KHz. All animals were left in the arena for at least 1 hour before recordings started in order to allow isoflurane to metabolize and the sedation effects to wash out. Sessions were always performed on the same hours of the day, during lights-on periods (when rats sleep the most), from 10 am to 4 pm (±30 mins), to maximize sleep time and to minimize variability related to the natural circadian rhythm of the animals (Figure 1C). To avoid excessive exploration and allow natural expression of the SWC, all animals were left to explore the arena for an additional 15 minutes after handling (see behavioral testing section), also for five consecutive days, twice a day, before surgery (Figure 1A).

### 2.5. Wideband signal pre-processing

Raw signals were fed to a MATLAB® based custom pre-processing pipeline developed in our lab (NigeLab, https://github.com/barbaLab/nigeLab). After conversion to MATLAB® compatible files, signals were digitally filtered in the local field potentials range (LFPs; < 300 Hz) and multi-unit activity (MUA; 300 – 3000 Hz) bands. LFPs were also down sampled to *f_s_ =* 1000 Hz. Both signals were then visually inspected and channels with excessive noise were masked out of the analysis. Furthermore, two animals, one of each group, had their electrophysiological recordings discarded as they were unrecoverably plagued by movement artifact or power grid noise. Thus, the sample number of electrophysiological results is N = 6 for CTRL and N = 5 for LESION groups.

### 2.6. Analysis of LFP signals

LFP signals were processed along two main lines of analysis: (1) *sleep architecture*, which includes global forebrain dynamics based on spectral features and the staging of the SWC as a sequence of stable states (i.e., hypnogram); and (2) *detection of specific sleep events*, particularly sleep spindles and slow oscillations. For sleep architecture, we first generated bi-dimensional state maps of spectral content following methodology originally described elsewhere (Gervasoni et al. 2004). Briefly, spectral decomposition of the local field potential (LFP) was performed independently for each recording channel using short-time Fourier transform for power spectral density estimation with LFP signals being segmented into 2-second windows with a 50% overlap. From the resulting time-resolved PSD *P*(*f*, *t*), two empirically defined spectral power ratios, i.e. Ratio 1 (cf. Eq. 2) and Ratio 2 (cf. Eq. 3) were computed for each channel at each time point *t* (frequency intervals in Hz):

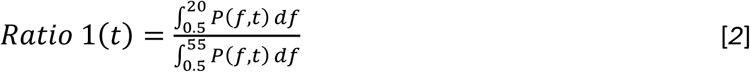

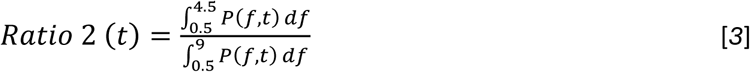

Principal component analysis (PCA) was then applied channel-wise and the so obtained first PCs for each ratio, smoothed using a 20-second Hanning window, were used to construct a two-dimensional state space. This representation consistently revealed the emergence of distinct clusters of points in the spectral state space. Clusters were objectively identified using non-overlapping isometric contours of an enhanced density map, which is constructed by dividing the density of points (estimated on a discrete grid) by the square of their instantaneous transition velocity. This procedure highlights regions of the state space that are both spectrally similar and temporally stable, effectively delineating discrete states. As previously shown, such clusters reliably correspond to the three canonical stages of the sleep–wake cycle, i.e. WK, SWS, REM, and occupy stereotyped locations in the spectral state map, allowing for objective sleep staging and construction of hypnograms. Points falling outside of these clusters were defined as belonging to the IS state, a transitional stage known to mediate important changes particularly between SWS and REM (see e.g. (Blanco et al. 2015)). Notably, previous work has also demonstrated that while the SWC architecture remains stable in healthy animals, it undergoes reproducible changes in response to brain perturbations (see, for example, (Dzirasa et al. 2006)). To ensure consistent and directly comparable results across animals when comparing geometric features of the SWC architecture, a multi-animal state space was also computed by performing PCA on the pooled ratio data from all subjects. Single subject tagged data was then projected onto the shared PCA basis, thereby avoiding inter-subject variability in the axes. Parameters such as area and centroid position of the SWC clusters in the shared space were then extracted and directly compared between the experimental groups. Although previous work has shown that this state-space framework can effectively identify REM sleep, particularly when supported by recordings from hippocampus, this stage was not included in the present analysis due to the absence of recordings of such brain region.

### 2.7. Detection of relevant sleep events

LFP was also processed for the detection of sleep spindles and Slow Oscillations (SOs). Spindles are ‘waxing and waning’ 10-15 Hz waves with a duration of at least 0.5 s (Blanco-Duque et al. 2024). SO are transient isolated oscillations characterized by a pairing of up and down states that recur at low frequency, with spectral content in the delta band (< 4 Hz) (Steriade et al. 1993). In both cases, we used a modified version of the well-established MATLAB® routine developed and made publicly available elsewhere (Kim et al. 2022). Briefly, spindles detection is carried out by first filtering in the interest band using a phase-corrected 6^th^ order high pass Butterworth FIR filter of 10 Hz cutoff frequency and of 8^th^ order low pass at 15 Hz. The envelope is then computed by using a Hilbert transform, which is convoluted with a 500 ms gaussian wave. Spindles were detected when the resulting signal exceeded 2.5 times the standard deviation. By computing the distribution of all intervals between start and peak of spindles and selecting those that better fit to the description, we carried out a data driven curation of such events. By its turn, for the detection of SO, the LFP signal was low-pass filtered at 4 Hz. During NREM sleep, all positive-to-negative zero crossings were identified along with preceding peaks and following troughs; up- and down-states were defined using the top 15% of peaks and bottom 40% of throughs, respectively. Differently from past studies, here we identified spindles and SO on a per-channel basis.

Once detected, the following metrics related to these electrographic events were computed: number of occurrences, duration, distribution of delays between SO and closest spindles, and wavelet power of spindles. Comparisons between groups were carried out in a stage- and area-specific manner. Moreover, channels were also rejected based on outlying spindles count, using standard Tukey’s fence approach (see section 2.10).

To inspect possible interactions of the ischemic lesion with the fundamental structure of spindles during SWS and IS and, given the precise temporal and spectral characteristics of such events, we computed metrics related to the time-frequency domain. For each animal and for both sleep states, 1.5 s of LFP activity was extracted from the longest (i.e., the most representative) bout, by centering the temporal window on the peak of each spindle. Then, given its widely proven efficacy in describing non-stationary oscillatory events (Sitnikova et al. 2014; Adeli et al. 2003), the Continuous Wavelet Transform (CWT) was applied to the segments to obtain spectrograms *S*(*f, t*). Briefly, CWT consists of the convolution of signal *x(t)* and complex Morlet wavelet as described in Eq. 4; *ψ* is the mother Morlet wavelet, *s* represents the time scale, while *τ* is a time-shift. Note that, Ω = 2*π* optimizes the spatio-temporal resolution (Pavlov et a., 2012) and results in the simple relationship between time scales and frequencies *f* ≈ 1/*s*. CWT was computed using MATLAB® 2024B’s `cwtfilterbank` function (with the ‘`amor`’ analytic Morlet wavelet) in the interval 10 – 15 Hz, consistently with spindles detection, and `wt` function to compute the complex wavelet coefficients *W*(*s*, *t*) and then *S*(*f*, *t*) (Eq. 4).

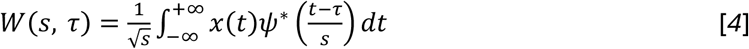

where

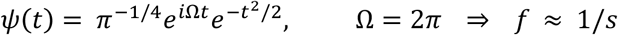

The central frequency, *f_central_*, represents the frequency at which the time-averaged spectrogram expresses the highest power (Eq. 5). On the other hand, the mean frequency, *f_mean_*, is the time-averaged instantaneous frequency, *f_inst_* (i.e., the frequency at which the power is maximum at a given time point - Eq. 6 and 7, respectively). Despite them being similar concepts, these two metrics describe differently the “spectral signature” of spindles in relation to their non-stationary (i.e., dynamic) nature. *N* stands for the total number of timepoints in the inspection window (i.e., 1.5 s × *f_s_* = 1500 samples).

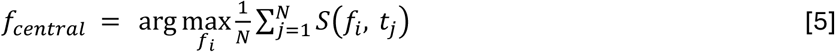

where

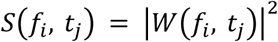

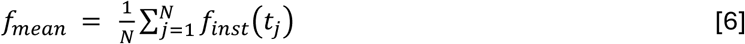

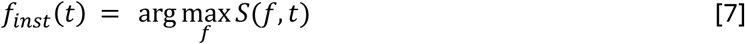

### 2.8. Detection of spikes and analysis of Single Unit Activity

With the aim of investigating the activity at the level of single neuronal units, we adopted the SWTTEO (Stationary Wavelet Transform and Teager Energy Operator) method for the detection of neuronal spikes (Lieb et al. 2017). Spike sorting was performed using the freely available superparamagnetic clustering technique developed by the Quiroga group (Quiroga et al. 2004). The resulting single-unit activity (SUA) was then analyzed in its average level and temporal structure of firing with two widely acknowledged metrics, the mean firing rate (MFR) and the Local variation Revised (LvR), which have been used in previous studies(Averna et al. 2020; 2021; Carè et al. 2022). MFR is simply computed as the average number of spike events over a temporal interval. Moreover, MFR was used to define a data-driven ‘bad unit’ rejection based on CTRL group to avoid including any bias introduced by the lesion induction at this level. A minimum MFR value, corresponding to the 10^th^ percentile of the whole-recording MFR distribution in the control group, was set to exclude poorly detected neurons. LvR considers the distribution of the inter-spike interval (ISI) but, unlike other metrics often used in this context, it is invariant to fluctuations of the firing rate. Indeed, by considering the instantaneous variability of the firing rate and the dependency of the ISI distribution on the refractory period following a spike, LvR holds three majors features: (1) firing irregularity is computed independently from the firing rate, (2) non-stationarity of the signal is considered by normalizing ISIs by the instantaneous firing rate (IFR), and (3) it encodes the distance of ISI distribution from a Poisson distribution as the deviation from LvR = 1, with lower values indicating a more regular (i.e., tonic) activity and higher values a more ‘burstier’ (i.e., clustered-in-time) firing. Given *I_i_* the *i*^th^ ISI, *n* the total number of intervals, and *R* the refractoriness constant, set to 5 ms according to previous literature (Averna et al. 2020), LvR is computed as in Eq. 8 (see (Shinomoto et al. 2009) for a thorough insight about the metric definition).

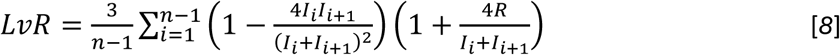

Both the metrics have been computed for all the retained units in a stage-specific fashion, separately for WK, SWS, and IS. Only stage bouts longer than 3 seconds were considered. For each unit, the metric average was calculated as a weighted average across the valid bouts, using their durations as weights. Moreover, to avoid confounding effects inherent to stage transitions, the first and last seconds of all viable bouts were discarded.

### 2.9. Peri-Spindle time histograms (pS)

In order to understand how the neuronal MUA, either underlying or induced by network events crucial for sleep-related functional reshaping, is affected by the lesion, the peri-spindle time histograms (pS) were computed. The aim of this approach was to target precisely the interplay of events at disctinct spatio-temporal scales, i.e., the micro-to-meso coordination of brain activity. For this specific analysis, the firing activity was considered on a per channel-basis given that spindles were also detect channel-by-channel (section 2.7). Similarly to the processing pipeline adopted for SUA, channels of MUA were data-drivingly retained according to a whole recording MFR threshold. The latter was defined as mean minus standard deviation, computed over the distribution of CTRL group’s channels to avoid any possible bias resulting from the lesion. Coherently to SUA analyses, sleep bouts shorter than 3 sec were discarded. pS was defined as the count of action potentials, in the same recording channel, in a 2 sec long window centered on spindle peak and binned over intervals of 10 ms. Histograms were computed separately for SWS and IS, in both S1 and RFA. Each good channel was scanned for spindles and an average pS was produced. Here, since the average level of activity was already addressed by the study of MFR in SUA, z-score normalization was provided as described in Eq. 9:

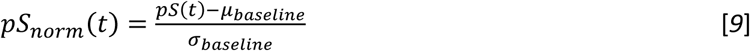

Channel’s firing rate within the stage of interest was first binned (10 ms bins) and then its average *μ_baseline_* and standard deviation *σ_baseline_* were estimated to obtain the normalized histogram *pS_norm_*(*t*). The animal’s representative histogram was then computed by averaging channels results. In the end, we extracted average and maximum values to test CTRL and LESION groups for any significant difference. Note that the average is equivalent except for a constant proportional to the area under curve, which is commonly adopted in the context of peri-event time histograms.

### 2.10. Statistical analysis

In all analyses, two-sided tests were conducted with a nominal α-level of 0.05. Continuous variables were first screened for outliers, defined as observations lying beyond the first or third quartile ± 1.5 × the interquartile range, and such values were excluded from inter-group and inter-stage comparisons. For simple, nonparametric pairwise comparisons, the Mann–Whitney U test was applied (p < 0.05) in accordance with our pre-specified sample plan (approved ethical protocol). To control the false discovery rate across multiple hypotheses, we applied the Benjamini–Hochberg procedure at a 5% threshold (Benjamini & Hochberg, 1995).

When hierarchical or repeated-measures structures were present, we fitted linear mixed-effects models (LMMs) or generalized LMMs (GLMMs) to account for within-subject correlation, improve power, and capture individual variability. All models were estimated in R (version 4.4.3) using the lme4 package (Bates et al. 2015). Fixed effects and planned contrasts were tested on estimated marginal means via the emmeans package (Lenth et al. 2025), with post-hoc multiplicity again controlled by Benjamini–Hochberg FDR (q < 0.05). Model fit and assumptions were evaluated both quantitatively and graphically using the performance package (Lüdecke et al. 2021).

The primary mixed-effects models took the following forms:

- Latency to first bout, log transformed LMM

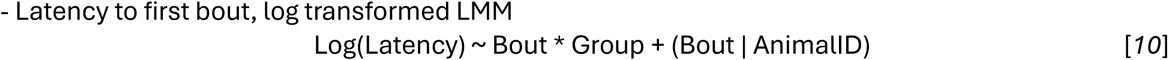

- Sleep-stage centroid analysis, LMM

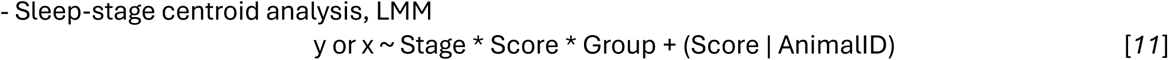

- Spindle centroid in sleep stage, LMM

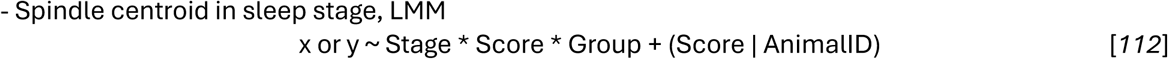

- Spindle spectral content, log transformed LMM

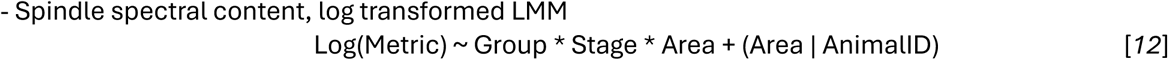

- Multi-unit activity, GLMM, Gamma family with logarithmic link function

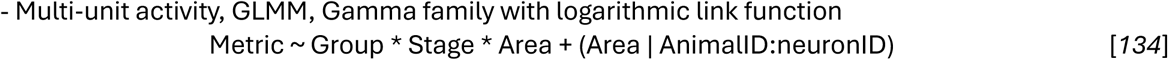

- Peri-spindle time histogram, LMM

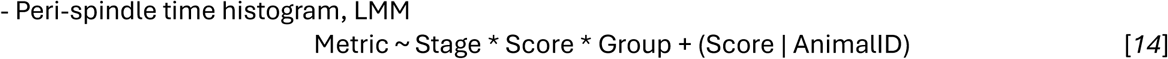

The magnitude of the observed effect (i.e., effect size) was computed using the conditional Cohen’s *d* for simple pairwise contrasts, and on marginalized means when LMMs were adopted. Note that, even if in the case of log-transformed LMMs *d* is computed over the transformed domain, it still holds the same meaning. *R_m_*^2^ and *R_c_* ^2^ were computed to have an insight into the variance explained by fixed effects and by the model as a whole, respectively, and to provide an alternative to *d* in the case of GLMM modeled data (Nakagawa et al. 2017). All the previously cited metrics and diagnostic plots are reported separately for each LMM or GLMM model (Supplementary material).

## 3. Results

### 3.1. Ischemic lesion induces motor dysfunction in the impaired limb

Animals that underwent ischemic lesion by microinjection of ET-1 in the motor area (CFA) clearly displayed alterations in the motor function which were indicative of impairment and, thus, efficacy of the model. In the GWT, the LESION group displayed an increased number of faults with the right limbs, which were contralateral to the lesion, in comparison to CTRL. This effect was particularly strong with the forelimb (**p = 0.002; Figure 1D, right top panel) and was also present in the hindlimb (*p = 0.037; Figure 1D, right bottom panel). There were no differences with ipsilateral limbs (Figure 1D, left panels). Additionally, in the CMT, there was a clear preference for using the intact left paw for supporting the body weight on the cylinder wall during rearing. This is shown in the increased number of touches of the wall with the left paw by animals of the LESION group (**p = 0.006; Figure 1E, left top panel) when compared to CTRL. Analogously, LESION animals displayed and increased value of the asymmetry index, indicative of a left paw preference (*p = 0.037; Figure 1E, right bottom panel). There was no such effect when animals used the right paw-only or both paws simultaneously (Figure 1E, top right panel and bottom left panel).

### 3.2. The ischemic lesion alters sleep dynamics and behavioral correlations at the macro-architectural level

The effects of the lesion in the global forebrain dynamic across the SWC are shown in Figure 2. Figure 2A shows representative hypnograms for every animal, while Figure 2B displays the distribution of latency (from the beginning of the recording session) to the occurrence of the first SWS bout lasting at least 30 s (top row) or at least 120 s (bottom row). Animals in the LESION group displayed significantly shorter latencies in both cases (30 s, CTRL/LESION = 3.17, p = 0.01; 120 s, CTRL/LESION = 2.24, p= 0.004; other latency values were omitted here for simplicity but are reported in Supplementary Figure S1). We did not observe a significant difference between groups regarding the duration of the classified SWC stages, although a trend to decreased WK and in increased SWS was noteworthy (Supplementary Figure S2, panel A, top row). Animals of both groups displayed clearly separable stable states in their stereotypical positions (Figure 2C, main panel) with the distribution of points in the SWS cluster of LESION animals assuming a bimodal shape, instead of the usual unimodal bell-shaped distribution of other clusters (Figure 2C, top and right panels). Yet, there were no significant differences in the size (area) of the clusters in the state map (Supplementary Figure S2, panel A, bottom row). Furthermore, LMMs showed significant patterns of correlation between the y ordinate (Ratio 2) of the clusters in the state map and the asymmetry index, computed for the CMT for assessing the motor performances (Figure 2D). While in the CTRL group there was no correlation and slopes resulted close to zero (slope for WK: 0.11 and for SWS: -0.15) in both the investigated states (WK and SWS), LESION animals displayed significant correlations. Specifically, WK tended to be negatively correlated (slope: -0.33; CTRL – LESION p = 0.15) and SWS showed a positive correlation (slope: 0.69; CTRL – LESION p = 0.02), demonstrating significant divergence between groups.

**Fig. 2.**
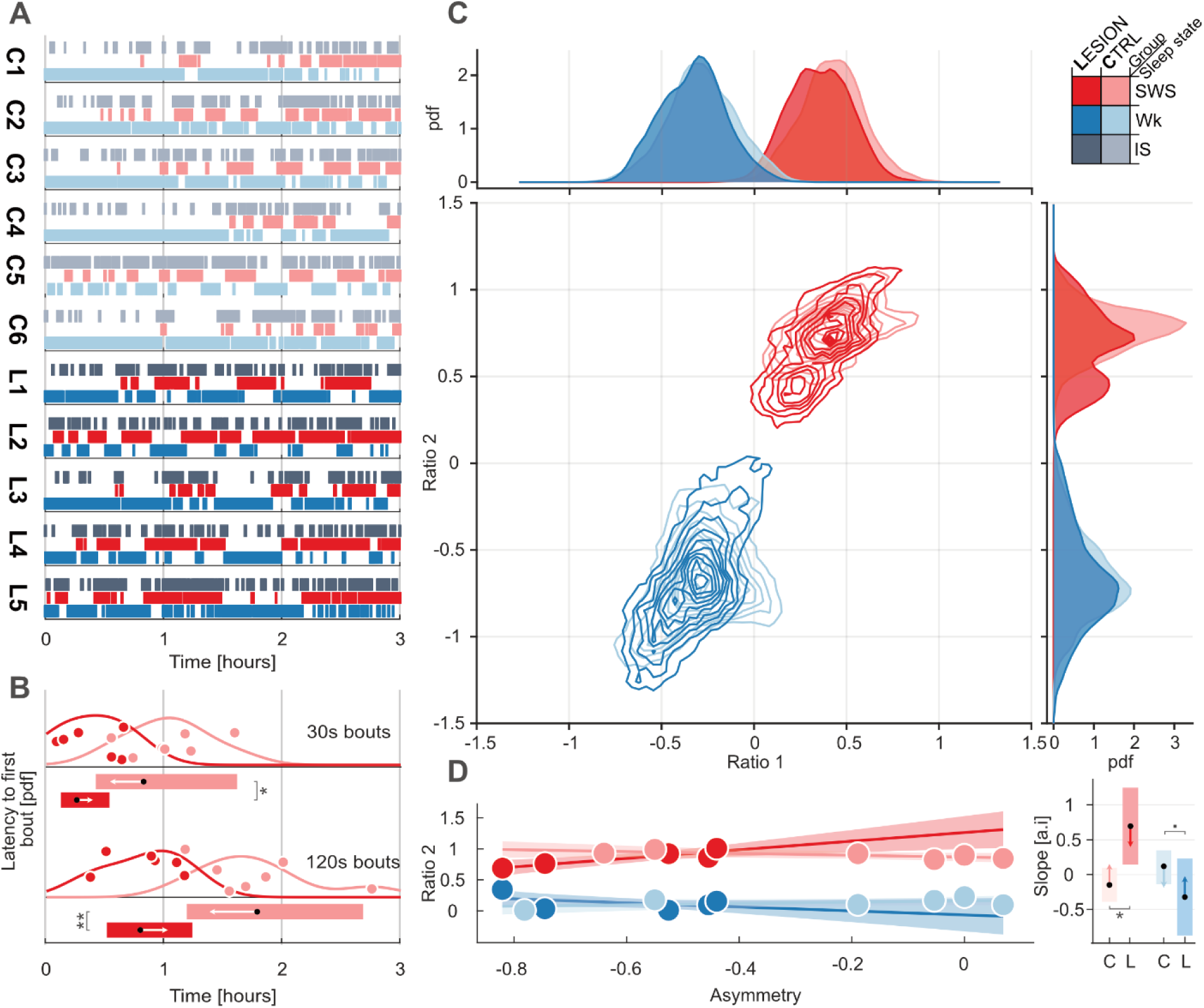
Macro-architecture alterations of the sleep–wake cycle. (**A**) Representative 3-hour hypnogram illustrating fragmentation across the sleep-wake cycle (SWC) in control (C1–C6) and lesioned (L1–L5) animals. Wakefulness (WK, blue), Intermediate Sleep (IS, gray) and Slow Wave Sleep (SWS, red); saturations indicate group (desaturated for CTRL, saturated for LESION). (**B**) Latency to first sleep bout above 30 s and 120 s. Kernel densities estimations with overlaid individual animal datapoints are depicted in light red for CTRL and dark red for LESION. Estimated marginal means (black dots) with 95% confidence intervals (boxes) and corrected pairwise comparison intervals (arrows, computed via *emmeans* in R) are shown below (*p < 0.05, **p < 0.005, LMM BH-FDR corrected). (**C**) Sleep state probability density functions (above, right) and two-dimensional state space density contours (center) derived from the first two spectral ratios (Ratio 1, Ratio 2) as in Gervasoni et al., 2014. Marginal densities illustrate shifts in sleep architecture between CTRL (desaturated blue/red/gray) and LESION (saturated blue/red/gray) animals. (**D**) Linear mixed-effects model (LMM) estimates of the relationship between behavioral asymmetry index (normalized units) and the y component of the state space centroids (Ratio 2) for each animal. Group-level estimated regression lines (solid) and 95% confidence bands (shaded area) reffect desaturated (CTRL) versus saturated (LESION) hues. The insert graph on the right reports group-level mean slopes (black dot) with 95% CI (colored box) with multiplicity corrected comparison intervals (arrows) indicating significant divergence between the two groups (* p < 0.05, ▪ p < 0.1, LMM BH-FDR corrected).

### 3.3. The ischemic lesion disrupts the coordination of sleep electrographic events at the meso-scale level

Next, we investigated how the ischemic lesion impacted the generation and coordination of spindles and SOs, which are plasticity-related electrographic events of major importance. We found that the fine temporal coordination between these two events, crucial for brain function, was disrupted following the ischemic lesion. We first computed the delay between SOs and spindles: while negative values indicate SO preceding spindles (and possibly causing them), positive values represent the other way around, as depicted in Figure 3A. Figure 3B shows the normalized occurrence of delays between SOs and spindles in 11 different temporal bins ranging from -1.5 s to 1.5 s. We found that negative delay values show a clear peak (particularly for RFA results), while positive values are uniform overall, suggesting, as expected, that SO precedes (and causes) spindles. In RFA, there were no differences in the occurrences of any delay between events across the groups. Differently, in S1, while there was no difference in SWS, the LESION group showed significantly less temporal coordination in the -600 ms (±150 ms) range during IS (*p = 0.0476, Mann Whitney BH-FDR corrected) with respect to the CTRL group. Then, we assessed the spindle features. Regarding their occurrence in the SWC, animals of both groups displayed not well separable conditions for both SWS and IS spindles. As shown by their overlapped distributions (Figure 3C, marginal panels around the main one), no significant differences in the average position (centroid) of spindle occurrence on the state map were found between groups, even though the distribution of Ratio 2 during SWS assumed, also here, a bimodal shape. Similar to what was investigated in section 3.2, LMMs were employed to probe the correlation between the y ordinate (Ratio 2) of the spindle clusters in the state map and the asymmetry index, computed for the CMT (Figure 2D). As in the previous case, we found no correlation for the CTRL group (slopes close to zero, see below) in both the investigated states (IS and SWS), while LESION animals displayed significant positive correlations. SWS showed a very detectable correlation (slope: 1.26, CTRL – LESION = -1.57, p = 0.007, Figure 3C), while IS showed a milder one (slope: 0.85; CTRL – LESION = -1.1, p = 0.032, Figure 3C). Finally, we found no differences in the quantity or in the duration of electrographic events (i.e. both SOs and spindles) across any brain regions, nor in any of the sleep phases during which they occurred (supplementary Figure S3).

**Fig. 3.**
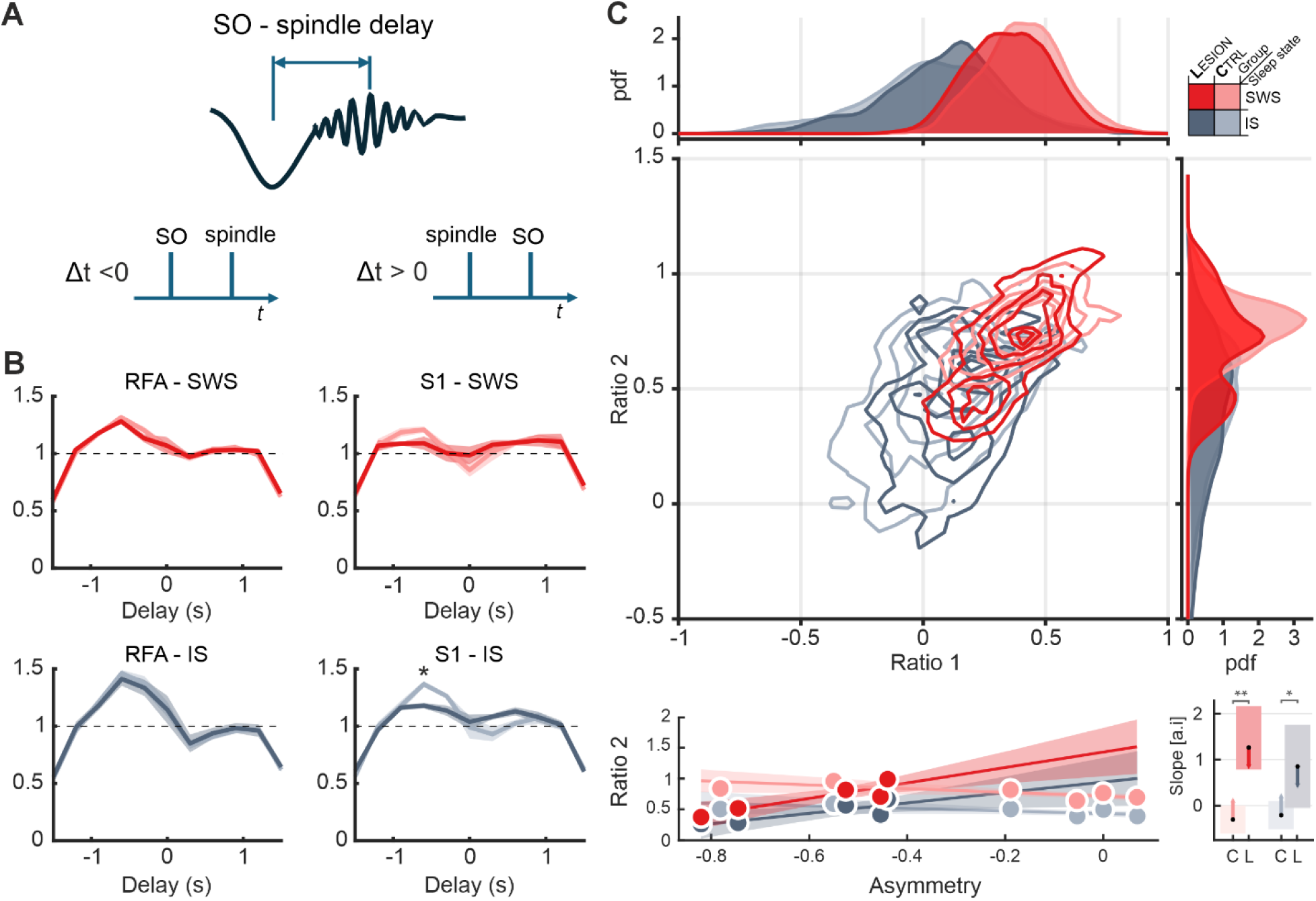
Meso-architecture alterations of network sleep events. After detection of spindle and SO events, the distribution of delays between them was computed and compared between groups, and their positions in the state map were assessed. (**A**) Delays were measured as the time interval from the throughs of SO and the peak of the nearest spindles and were binned into 11 central values -1.5 s to 1.5 s (300 ms separation from one another), with negative values meaning SO precedes spindles and positive values the other way around. (**B**) Normalized distribution of delays for RFA (left panels) and S1 (right panels) during SWS (top panels) and IS (bottom panels) are shown, with unsaturated colors for the CRTL group and saturated colors for the LESION group. While in all cases a reasonably consistent relationship of SO preceding spindles is shown, animals of the LESION group show a decreased relative number of 600 ms (± 150 ms) delays in S1 during IS (* p = 0.047, Mann Whitney U test, BH-FDR corrected). (**C**) Spectrally determined position of SWS and IS spindles in the state map (main central panel). Their centroids (mean position of spindles computed for each state) did not differ between groups (distribution on the marginal panels around the central panel). Bottom panels show that Ratio 2 of the centroids correlate positively with asymmetry index in LESION animals only (**p < 0.01 during SWS, *p < 0.05 during IS; LMM, BH-FDR corrected). In all panels SWS is represented in red while IS in grey, with unsaturated colors for CTRL animals and saturated colors for LESION animals.

In order to further investigate the ongoing electrophysiological activity during spindles, their spectral content was assessed. Although total spindle power did not reveal any significant difference between groups, metrics related to the time-frequency decomposition captured lesion-induced changes. The average CWT time-frequency decomposition of spindle events during the most representative (i.e., longest) SWS and IS intervals was computed using 1.5 s windows centered on spindle peaks in both S1 and RFA. Results for LESION and CTRL groups and all the combinations of sleep stage and cortex area are shown in Supplementary Figure S4. The differential maps were computed by subtracting the spectrograms of the CTRL group from those of the LESION group. As qualitatively shown by differential maps in Fig. 4 (panels A1, A2 - referred to S1, during SWS and IS, respectively; panels B1, B2 – referred to RFA, during SWS and IS, respectively), spindles spectral power appears to be differently distributed in the time-frequency domain between CTRL and LESION groups, particularly in S1. Fig. 4C reports the CWT spectrogram of a representative spindle occurred during SWS of a randomly selected animal. Furthermore, it provides a graphical intuition about the computation of *f_central_* and *f_mean_* as previously described (cf. section 2.7). IS spindles exhibited lower *f_central_* and *f_mean_* in LESION animals compared to CTRL ones (inter-group comparisons, LMM BH-FDR corrected. *f_central_*: IS/S1 CTRL – LESION, p = 0.019; *f_mean_*: IS/S1 CTRL – LESION, p = 0.0016). A similar trend, despite not being significant, was observed for all the remaining combinations of sleep stage and implantation site (i.e., S1/SWS, RFA/SWS, and RFA/IS). Noteworthy, *f_mean_* variations across stages showed a group-dependent trend in S1, reflecting the effect of the lesion (Fig. 4, panel E. Second-level contrasts – difference-of-differences between matched stages: •p = 0.093, LMM BH-FDR corrected).

**Fig. 4.**
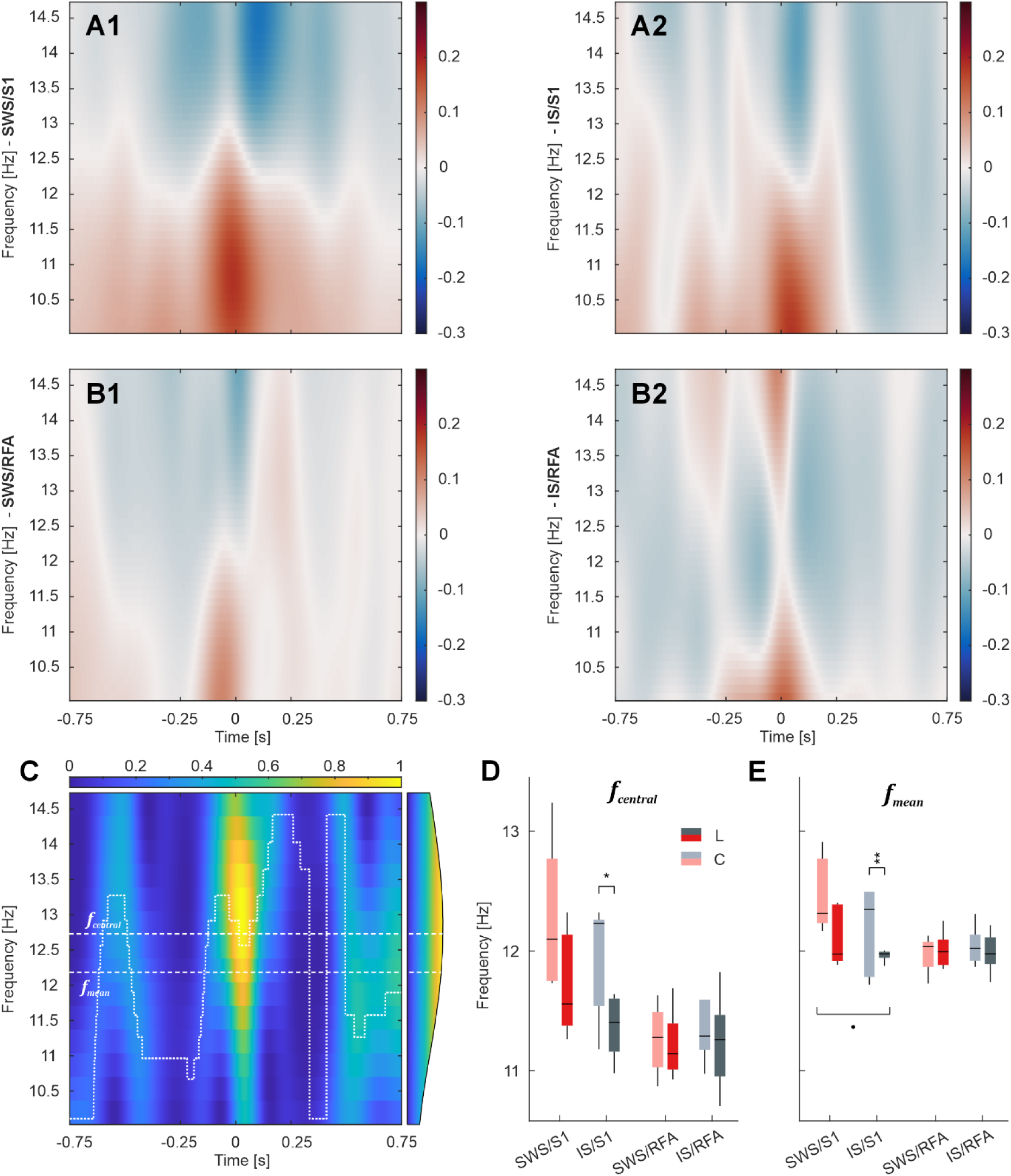
Spindle spectral content reflects stage-specific effects of the ischemic lesion. Group-average Continuous Wavelet Transform (CWT) spectrograms of spindles activity in the LFPs were computed separately for Slow Wave Sleep (SWS) and Intermediate Sleep (IS), and implantation sites, i.e., primary somatosensory area (S1) and Rostral Forelimb Area (RFA) (Supplementary Figure S4). **A1, A2)** Differential spectrograms obtained as the subtraction LESION – CTRL in S1 for SWS and IS (A1 and A2, respectively). Differences between groups are shown with their polarity via colormap, with red and blue representing positive and negative differences respectively. Maps were interpolated (bicubic spline interpolation) for visualization only. **B1, B2)** Differential spectrograms obtained as the subtraction LESION – CTRL in S1 for SWS and IS (B1 and B2, respectively), analogously to panels A1 and A2. **C)** CWT spectrogram of a representative spindle event occurred during the longest SWS bout of a randomly selected animal (C1, see Fig. 2 panel A). Normalized power scale around spindle’s peak (0 s timestamp, inspection interval [-0.75; +0.75]) is shown through a colormap. On the right, the time-averaged power spectrum used to compute spindle central frequency (*f_central_*), shown by a dashed line. A dotted line depicts the time evolution of the instantaneous frequency (*f_inst_*) inside the interval whose mean value is the spindle mean frequency (*f_mean_*), shown by a dashed line. **D)** *f_central_* distribution in CTRL and LESION (light and dark colors, respectively) for in S1 and RFA, during longest bouts of SWS and IS (red and gray, respectively). Results are shown as a box plot depicting median, first (Q1) and third quartile (Q3), whiskers (Q1 - 1.5 * interquantile range (IQR) and Q3 + 1.5 * IQR), and outliers. Distributions were tested for any inter-group (fixed site) and inter-stage (fixed group) statistically significant difference (inter-group: *p < 0.05 - LLM BH-FDR corrected; second-level contrasts – difference-of-differences between matched stages: °p = 0.067 - LMM BH-FDR corrected). **E)** *f_mean_* distribution, computed, shown and tested analogously to central frequency (inter-group: **p < 0.005 - LLM BH-FDR corrected; second-level contrasts – difference-of-differences between matched stages: •p < 0.1 - LMM BH-FDR corrected).

### 3.4. SWC differentially modulates the spiking activity at the *micro-architecture* level

Single-unit activity (SUA) patterns during SWS, IS, and WK in S1 and RFA are illustrated in Figure 5, where the qualitative evaluation of the raster plots confirmed that firing rates remained within physiological levels throughout the SWC (Figure 5 A1 for S1 and Figure 5 B1 for RFA). The quantitative investigation of SUA was conducted through the analysis of MFR and LvR (*cf.* section 2.8). The scatter plots of Figure 5 A2 (S1) and B2 (RFA) show the relationship between the firing rate of individual units during WK (x-axis) and SWS (y-axis) for both CTRL (blue dots) and LESION (yellow dots) groups. In S1 (Figure 5 A2), the points and corresponding histograms largely overlapped. Quantification of MFR confirmed the absence of stage-dependent changes in S1 for both CTRL and LESION animals (Figure A3). Differently, in RFA and only in LESION animals, a significant difference is observed between WK and SWS, with a trend between IS and SWS (inter-stage contrasts, GLMM BH-FDR corrected: SWS – IS RFA LESION °p = 0.067; SWS – WK RFA LESION p = 0.00038, Figure 5 B3).

**Fig. 5.**
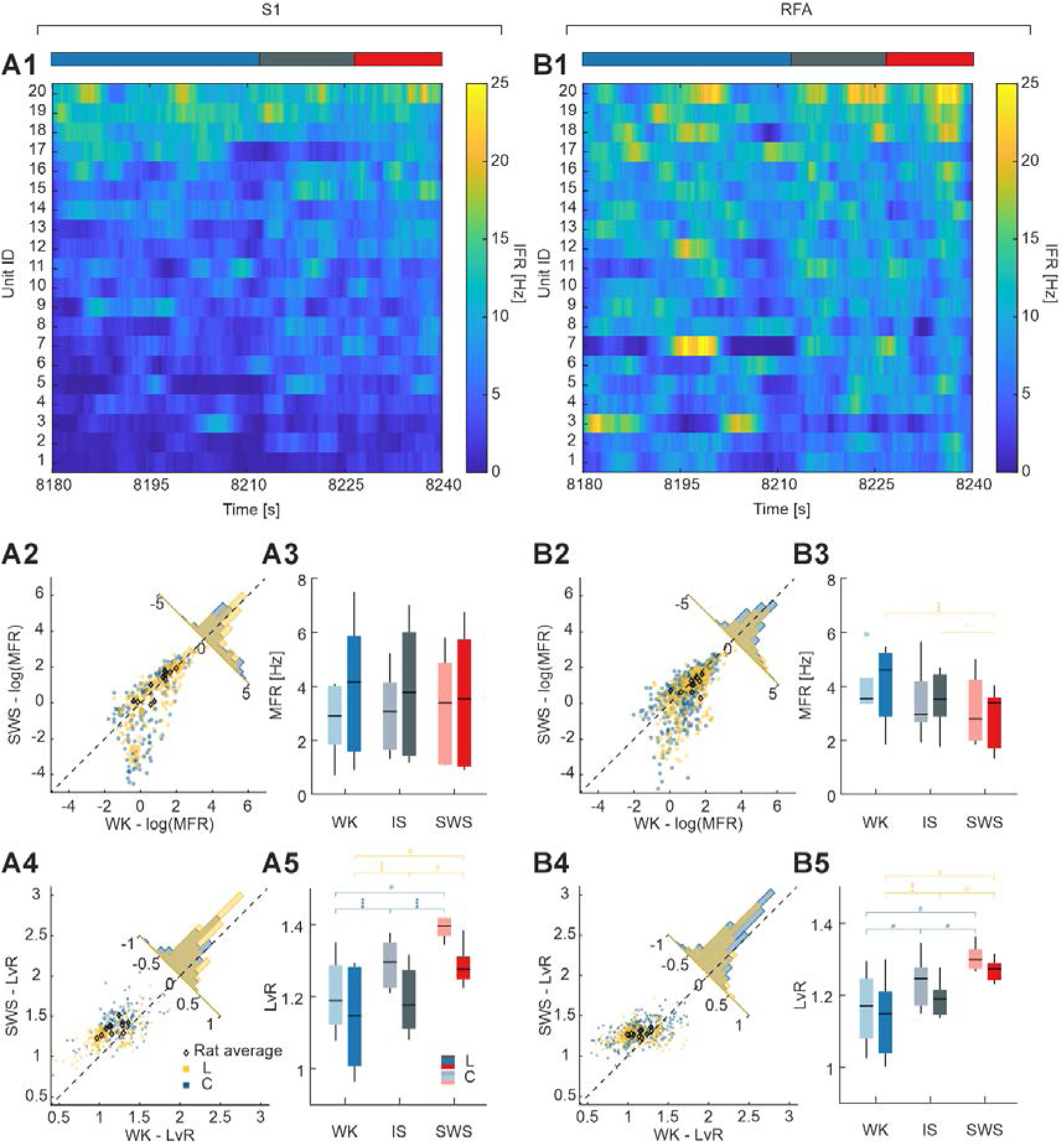
Micro-architecture assessment of stage-specific neuronal activity. Characterization of Single-Unit Activity (SUA) in key areas of the sensorimotor loop (S1 and RFA) across the SWC. S1 and RFA results are shown in panels A and B, respectively. **A1, B1)** Raster plots of the 20 most active units in a representative interval showing transition from wakefulness (WK, red) to Slow-Wave Sleep (SWS, blue) through Intermediate Sleep (IS, gray), in a randomly chosen animal (i.e. C6). The Instantaneous Firing Rate (IFR) is color-coded (blue to yellow as the firing rate increases). **A2, B2)** Distribution of units in relation to their Mean Firing Rate (MFR) during WK and SWS. Each dot represents a neuronal unit, while diamonds stand for animal averages (blue and yellow for CTRL and LESION groups, respectively). The corner histogram provides insights on stages equilibrium by projecting the scatter plot along the bisector of the axes. **A3, B3)** Animal average MFR distribution in CTRL (dark colors) and LESION (light colors) groups during WK, IS, and SWS (blue, gray, and red, respectively from left to right). Results are shown as a box plot depicting median, first (Q1) and third (Q3) quartile, whiskers (Q1 - 1.5 * interquantile range (IQR) and Q3 + 1.5 * IQR), and outliers. Distributions were tested for any inter-group (fixed site) and inter-stage (fixed group) statistically significant difference (inter-stage: °p = 0.067, ***p < 0.0005 - GLMM BH-FDR corrected; yellow marks refer to LESION group contrasts). **A4, B4)** Distribution of units in relation to their Local variation Revised (LvR) during WK and SWS. Results are shown analogously to panels A2 and B2. **A5, B5)** Animal average LvR distribution in CTRL and LESION group during WK, IS, and SWS. Results are shown and tested analogously to panels A3 and B3 (inter-stage: ***p < 0.0005; #p ∼ 0 – GLMM BH-FDR corrected; yellow and blue marks refer to LESION and CTRL groups, respectively).

The scatter plots of LvR (Figure 5 A4 for S1 and B4 for RFA) indicate highly similar distributions between WK and SWS in both CTRL and LESION groups. Interestingly, in both areas, the centroids of the LvR distributions during WK and SWS lie above the bisector line, indicating that both CTRL and LESION animals exhibit higher LvR values in SWS than in WK. This suggests that neuronal firing is more ‘burst-like’ during SWS in both regions. Interestingly, while during WK no difference is observed between CTRL and LESION groups, during IS and especially SWS there is a clear trend, even if not significant, between the two groups, both in S1 and RFA. Additionally, all the inter-stage comparisons both in CTRL and LESION resulted to be statistically significant (inter-group contrasts, GLMM BH-FDR corrected: CTRL/S1 WK – IS p = 0.44·10^-4^; CTRL/S1 IS – SWS p = 0.27·10^-6^; LESION/S1 WK – IS p = 0.00034; LESION/RFA p = 0.13·10^-6^; remaining contrasts p ∼ 0) (Figure 5, A5 for S1 and B5 for RFA).

Finally, the coordination between spindle events and MUA (i.e., micro-to-meso coordination) was addressed. Group average normalized peri-spindle time histograms, *pS_norm_*, clearly display a coordination of spindle events and MUA, consistent with the typical ‘waxing and waning’ shape of spindles, with sharp oscillations getting more and more prominent (i.e., deviated from *μ_baseline_*) approaching the spindle peak and decreasing symmetrically after the peak. While a similar waveform can be seen both during SWS and IS, either in S1 and RFA, the two areas of investigation display a slightly distinct pattern preserved along different sleep stages (Figure 6, panels A1, A2 – S1; B1, B2 – RFA). The average (Figure 6C) and maximum (Figure 6D) values of *pS_norm_* (within the interval [-0.75; +0.75] s from the spindle peak), representing the broad correlation between spindle and undergoing MUA and their mutual modulation strength respectively, do not show any significant inter-group or inter-stage difference (p > 0.05) in either S1 or RFA. Ultimately, a trend may indicate that the lesion affects the IS-SWS difference in CTRL and LESION in S1 (second-level contrasts – difference-of-differences between matched stages: (IS – SWS) S1 CTRL - (IS – SWS) S1 LESION, ° p = 0.067 - LMM BH-FDR corrected).

**Fig. 6.**
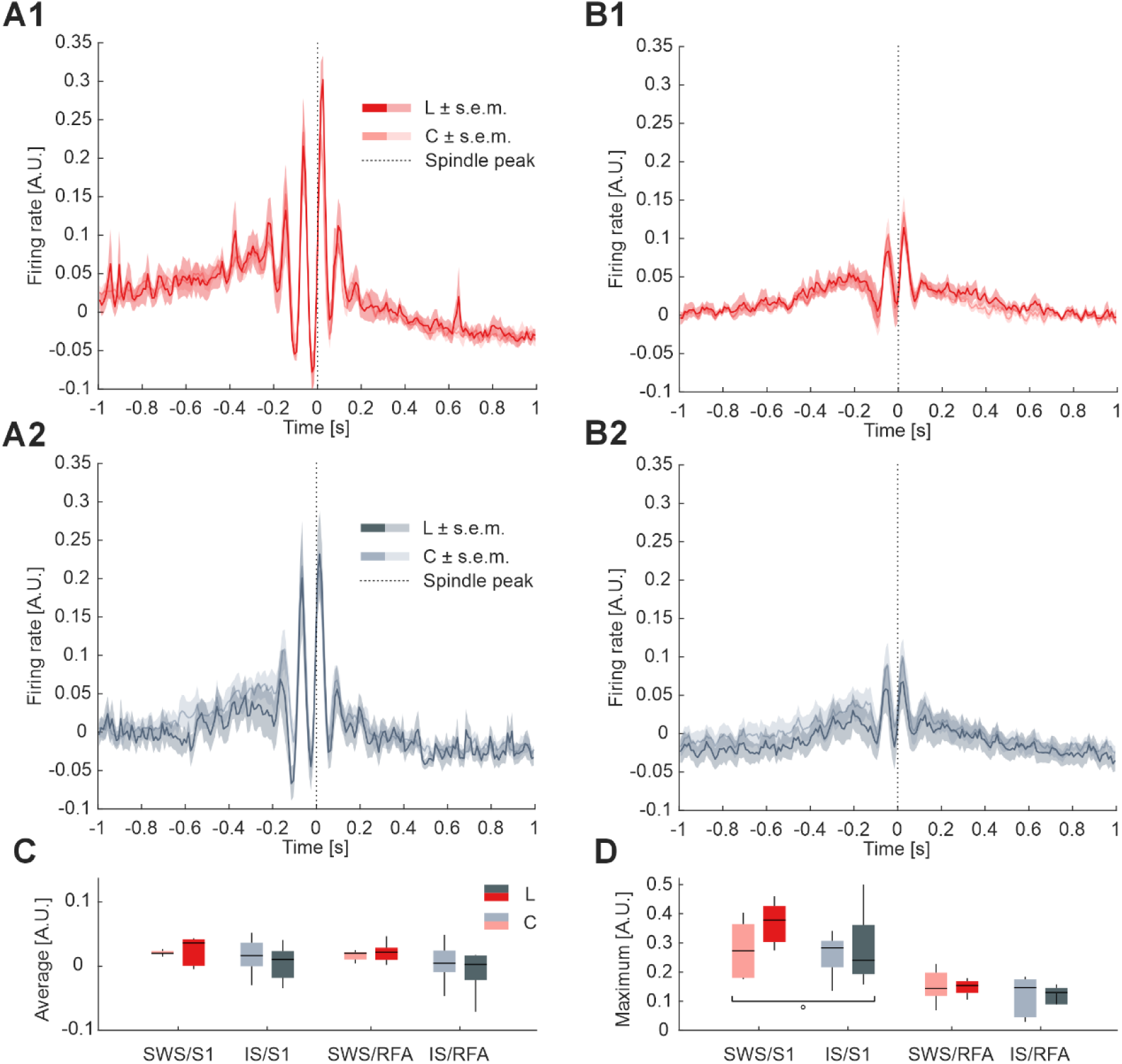
Micro-to-meso coordination of neural activities during sleep. Instantaneous Multi-Unit Activity (MUA) firing rate during sleep spindles, separately for Slow Wave Sleep (SWS) and Intermediate Sleep (IS), in primary somatosensory area (S1) and Rostral Forelimb Area (RFA). Panels A and B (portraying S1 and RFA, respectively) depict normalized peri-spindle time histograms (pS), z-score normalized, in CTRL and LESION groups. **A1, B1**) pS during SWS in CTRL (group mean ± Standard Error of the Mean – SEM; dark red line and shade) and (group mean ± SEM; light red line and shade). **A2, B2**) pS during IS in CTRL (group mean ± SEM; dark grey line and shade) and (group mean ± SEM; light grey line and shade). **C**) pS average distribution in CTRL (dark colors) and LESION (light colors) groups during SWS and IS (red and grey, respectively from left to right). Results are shown as a box plot depicting median, first (Q1) and third (Q3) quartile, whiskers (Q1 - 1.5 * interquantile range (IQR) and Q3 + 1.5 * IQR), and outliers. No statistical significance was found for any inter-group or inter-stage contrast. **D**) pS maximum distribution in CTRL and LESION group during SWS and IS. Results are shown and tested analogously to panel D (second-level contrasts – difference-of-differences between matched stages: ° p = 0.067 - LMM BH-FDR corrected).

## 4. Discussion

In this work we investigated the impact of an ischemic lesion of the primary motor cortex (CFA) in the spontaneous neural activity of the sensorimotor loop, specifically S1 and RFA. Using wideband electrophysiological recordings and comprehensive analysis along the several hours during the SWC, we not only assessed neural activity across different levels of brain organization (single unit, network, global neurodynamic, and behavior), but also the integration and coordination between the different scales, all in a stage-specific fashion. Behavioral data clearly showed robust motor deficit in animals of the LESION group (Figure 1D and 1E), which demonstrates that the ET-1 lesion model was effective. Considering the solid basis for the ET-1 model demonstrated in previous literature (Frost et al. 2006; Thaysen et al. 2024; Lin et al. 2022) and replicated here, electrophysiological findings will be anchored in the framework of a successful induction of cortical lesion in CFA. Furthermore, electrophysiological results showed a range of statistically significant modifications spanning from global forebrain dynamics to disruptions of network oscillatory events, while activity at the unit level displayed noteworthy trends. Two aspects of our current findings are of particular interest. First, instead of major electrographic changes such as of raw spectral content or decreases of unitary activity, LESION animals displayed rather substantial shifts on the fine tuning within and across the different scales, highlighting the importance of a multi-scale approach in studying sleep neural dynamics. Second, IS state, a brain activity-regime of major importance to neuroplasticity, seemed to be particularly affected.

Starting with the largest scale of brain organization, global forebrain dynamics were significantly impacted in LESION animals, including a decreased latency to the first SWS bout (Figure 2B) and a trend to an increased sleep duration (more SWS and less WK). The first two findings suggest that the lesion induces a sleep drive, an interpretation that aligns with prior literature showing elevated delta-band activity during wakefulness following cortical injury (Ahmed et al. 2011; Niedermeyer 1982; Kim et al. 2022; Duss et al. 2017). In the same line, in a recent study of our group with an overlapping dataset, we found increased values of interarea (S1 and RFA) synchronization in lower bands (delta and theta), even though no differences in the raw spectrum were found (Canu et al. 2025). Considering the delta-prevalent spectral signature of SWS, the well-established link between sleep and neuroplasticity (Vahdat et al. 2017; Brodt et al. 2023), the correlation between ischemic stroke and the induction of plastic susceptibility state by subproducts of neuroinflammation (Kane and Ward 2021), and the multiple benefits of sleep in brain recovery (Hodor et al. 2014), it is reasonable to interpret such results as evidence of a brain recovery mechanism by increasing SWS. On the other hand, some studies reported that ischemic stroke in humans induces longer latencies to sleep (Sterr et al. 2018) with longer periods of time spent awake (Suh et al. 2014). Similar findings were found in animals that underwent stroke produced by the occlusion of the middle cerebral artery; MCAO model (Sharma et al. 2022; Chu et al. 2025). It is not possible to derive, from the present data, the reason for this discrepancy. Yet, a natural interpretation is that both outcomes coexist and are separated in the space of possibilities by factors like the species (rats versus humans or mice), time of assessment (subacute versus acute), the experimental model (ET-1 versus MCAO or naturally occurring stroke), and particularly age. Regarding this last aspect, it is noteworthy the observation that the direction of sleep modulation (increases versus decreases) induced by stroke may get inverted when young animals or aged ones are compared to control animals (Chu et al. 2025).

When global forebrain dynamics were associated with motor behavior, we found that LESION significantly induced correlations between spectral content (ratio 2 of the state space) and asymmetry in paw usage (Figure 2D), significantly for SWS and a trend for WK. These correlations were absent (zero slope) in healthy CONTROL animals. This indicates that ischemic stroke alters the coupling between neural oscillatory patterns and motor performance. Furthermore, it supports the idea that SWC contributes actively to post-stroke circuitry (re)organizing. Or in physiological terms, the distinct plasticity mechanisms of the different SWC stages are recruited to promote brain rewiring reacting to the injury, putatively for motor function recovery. In fact, it is well known that the mnemonic role of SWC physiology is contingent on the presence of memory-encoding events (Ribeiro and Nicolelis 2004; Ribeiro et al. 2007), being absent if animals are not exposed to any novelty. In this context, the lesion can be understood as a strong plasticity-triggering event, consistent with previous observation related to neuroinflammation (Kane and Ward 2021). It is also noteworthy that the signal of the slope in SWS correlation is different from that of WK: while the first is positive the other tends to be negative. Given that left-paw asymmetry is coded as negative and that ratio 2 reflects delta dominance in the low-frequency band (Gervasoni et al. 2004), this implies that, in WK, delta power content is correlated to greater asymmetry, while in SWS it is correlated to less asymmetry. Such pattern suggests a maladaptive role of delta during WK and a beneficial, restorative one in SWS. In any case, these findings not only corroborate the critical role of sleep in brain plasticity and recovery following injury, but also highlight the differential and complementary roles of the different sleep stages in such processes.

From the network level, the impairment of the ischemic lesion is also evident, as can be seen from the disruption of the temporal coordination of key electrographic events, as well as from the different alterations on their characteristics, in animals of the LESION group. Overall, the distribution of delays between slow oscillations (SOs) and thalamocortical spindles (Figure 3B) were shown to be largely consistent with established neurophysiological mechanisms: SO up states facilitate the occurrence of spindles which is reflected as a peak in SO → spindle delays (negative; within the 450 to 750 ms bin) and flat distribution in spindle ◊ SO (positive) (Schreiner et al. 2021; Mölle et al. 2002). Yet, this pattern is significantly disrupted in the S1 area of LESION animals during IS, in which the peak disappears, and the distribution becomes approximately uniform. Furthermore, the average spindle was “slower”, in the LESION group, as both central and mean frequencies were shown to be decreased in those animals (Figure 4). Finally, there was a trend to smaller number of such events. In both cases, the difference was in S1 and during IS, only. This evident impact of the lesion on such important plasticity-related oscillatory activity is simultaneously very plausible and also provides novel insights on the consequences of an ischemic stroke. To begin with, it is very well known that the cortex, region target by the lesion here, has a central role in the generation of both SO and spindles. Briefly, while spindle are generated by a reciprocal excitation-inhibition interplay between neurons of the thalamus reticular nucleus (TRC) and thalamocortical neurons (TC), SO events are generated in reciprocal cortico-cortical projections and its up state provides excitation to corticothalamic projections that gate spindle events (Steriade et al. 1993; Steriade 2003). Following an ischemic lesion, corticothalamic projections may be impaired which would affect SO drive of spindles and, thus, their fine temporal coordination, which would be expressed as the flattening of the delay distribution. By the same principle, a cortical lesion would reduce the synchronization of spindles activity across the cortex, impairing its detection or slowing down its frequency content, very much in line with previous observation in the literature (Gottselig et al. 2002). Together with hippocampal ripples, SO and spindles act to promote consolidation of memories in distinct temporal and spatial scales (Staresina 2024). While SO clocks long-range interactions for content integration, spindles promote neuronal reorganization targeted especially in regions involved with the memory trace acquired during wakefulness. Ripples then tune neuronal firing rates to communicate encoding engrams. Above all, the correct temporal locking between SO and spindles helps promoting ripples (Helfrich et al. 2019), but also the efficiency of this triple coordination of oscillation in consolidating memory traces is largely determined by their correct nesting (Schreiner et al. 2021; Mikutta et al. 2019). Taken together under a single framework, the observed disruption of SO-spindle coordination would impact physiological sleep-dependent plasticity. On the other hand, putative increases of SWS (trend in this study) and isolated SO and delta waves (Kim et al. 2022), and possible neuroinflammation-induced plastic susceptibility, might promote maladaptive changes (e.g. post-stroke epileptiform activity). In other words, the physiological drive to sleep may be present, but the quality of the sleep itself, a separate concept, is lacking.

It is also noteworthy that lesion-induced modifications seen at the network level happened during IS, instead of SWS. Classically, major roles have been assigned mostly to canonical states of sleep: SWS as promoter of neuronal reverberation (Klinzing et al. 2019; Ribeiro et al. 2004) and REM as privileged window for neurochemical cascades underlying synaptic remodeling (Almeida-Filho et al. 2018). Yet, the transitions between states in general and IS in particular are not mere physiological noise or random useless activity, but instead, seem to play preponderant role as a bridge between memory reverberation and circuit remodeling functions. For instance, significant correlations between cortical spindle activity and hippocampal CaMKII phosphorylation (a major neurochemical biomarker of neuroplasticity) was observed only in animals that were not deprived of IS (Blanco et al. 2015). Of particular interest, in humans, sleep spindles during NREM stage 2 transiently synchronize hippocampo–striato–thalamo–cortical networks, providing a neural mechanism for large-scale coordination underlying motor memory consolidation (Boutin et al. 2018). Moreover, these spindles cluster in temporally organized bursts, and the strength of such clustering predicts overnight gains in motor performance (Boutin et al. 2024). In rodents, the IS sleep stage is the equivalent of human NREM-2 and, as previously mentioned, displays spindle-rich activity involved in memory consolidation.

When assessing the link between spindle activity and behavior, LESION animals displayed strong correlations, during both SWS and IS, between spectral content underlying the occurrence of spindles (spindle centroid in the state map; Figure 3C) and paw usage asymmetry in the CMT. In this case, both showed a statistically significant positive correlation, a relationship which was, instead, absent in CONTROL animals. Here, it is important to understand that the correlation is not with the spectral content of the spindles themselves, but actually of the background oscillation pattern during their occurrence. As before, increases of the delta activity occurring in the background of spindles predicts less asymmetry in paw usage during both IS and SWS. In line with the previous interpretation, these results point to a possible participation of spindles in the restorative effects of sleep following stroke.

Finally, in the investigation of SUA, the impact of the lesion was less apparent (Figure 5). Overall, there were no significant differences in the MFR or LvR between LESION and CONTROL groups, only across stages of the same group, such as a decrease of MFR in RFA from WK to SWS (Figure 5 B3), and an increase of LvR in both areas from WK to IS, and from IS to SWS (Figure 5 A5 B5). In fact, a burstier (increased LvR) activity during SWS was expected. Indeed, both SWS and IS physiologically exhibit slow wave activity and in particular SOs, which consist of alternance of network states of hyperpolarization (i.e., ‘down’ states – periods of general silencing of firing activity) and depolarization (i.e., ‘up’ states – periods of higher neuronal excitability), facilitated firing resulting in burst activity (Adamantidis et al. 2019). Despite LvR inter-group contrasts not being statistically significant and considering the higher variability of spiking activity with respect to macro- and meso-scale activity (fundamentally observed via LFP) (Logothetis 2003), trends are observable. Particularly in S1, and more during SWS and IS than in WK, SUA appears more regular (i.e., LvR reduction in LESION ◊ less‘bursty’ activity) (Figure 5 A5 B5). This could suggest a plausible effect of the lesion in the transmission of memory-encoding engrams, as revealed by their regularity. Future studies will be necessary to assess that changes in the neuronal firing regularity are related to the pathophysiology of stroke

The same applies to the assessment of the coordination between MUA and spindles made by pS (Figure 6); none of the inter-group contrasts were statistically significant. However, similarly to what was observed for *f_mccc_* in S1 (Figure 4E), second-level contrasts on the maximum modulation level indicate a trend of fragmentation in sleep stages balance following the ischemic lesion (Figure 6D). Altogether, the results show a higher sensitivity of S1 to lesion compared to RFA and point to an impairment of sleep continuity by compromising sensory gating during sleep with a possible interference with underlying sleep-dependent plasticity, as spindles are known to play a key role in sleep homeostatic maintenance (Urakami et al. 2012; Fernandez and Lüthi 2020). Notably, previous studies on both rodents and humans have found topography-specific patterns in spindle occurrence across cortical areas (Kim et al. 2015; Martin et al. 2013), similarly as here, once we observe a distinct pS shape depending on S1 and RFA. This set of results point to a greater resilience of activity in the cellular, micro-circuit level of brain organization and this is reasonable to a certain extent, considering that an ischemic stroke induces a circuit disconnection and not a dysfunction of the neuronal cell (Guggisberg et al. 2019; Grefkes and Fink 2014).

Overall, our set of results clearly show that an ischemic lesion has a considerable impact on neural activity in many levels of brain organization, being stronger at the network level, but also in global dynamic and behaviour, and considerably less at the micro-architecture. Such observations were all performed in regions that are not the direct target of the lesion, although having close functional relation to it as part of the sensorimotor cortical loop. This is strong evidence that surviving parts of the cortex clearly react to the lesion in a putative restauration reorganization effort. If this response is enough and to what degree it efficaciously promotes recovery is yet to be fully determined, but certainly it depends on the nature of the lesion and the affected area, as well as an array of external circumstances including, among others, age, gender, species (Roy-O’Reilly and McCullough 2018)(. Future studies may determine the exact contribution of each of the SWC alterations observed in these and other studies towards promoting plasticity and if they are beneficial or maladaptive. In any case, it is evident that this network reorganization process represents a privileged window for therapeutic intervention (Silasi and Murphy 2014) that can and should be leveraged also by neurotechnologies. For instance, by using responsive coordinated neurostimulation aimed at suppressing aberrant electrographic events (see for example: (Réboli et al. 2022)) and simultaneously at restoring healthy level of oscillatory synchronization of electrographic events (e.g. (Reinhart and Nguyen 2019)), one may be able to promote enduring rehabilitation in patients victims of ischemic stroke in a personalized form (Carè et al. 2024).

## Acknowledgments and Funding

The author(s) declare that financial support was received for the research and/or publication of this article. This research was supported by #NEXTGENERATIONEU (NGEU) and funded by the Ministry of University and Research (MUR), National Recovery and Resilience Plan (NRRP), project MNESYS (PE0000006)-A Multiscale integrated approach to the study of the nervous system in health and disease (DN. 1553 11.10.2022). This research has been also supported by the NATO Science for Peace and Security Programme (BIONIC, #G8447) to MC. VRC was initially supported by the Marie Skłodowska-Curie Individual Fellowship MoRPHEUS, Grant Agreement No. 101032054, funded by the European Union under the framework programme H2020-EU.1.3.-EXCELLENT SCIENCE.

