## Supplementary for "Multi-scale disruption of sleep-related cortical activity following ischemic lesions"

**This PDF file includes:**

Figs. S1 to S4

**Fig. S1.**

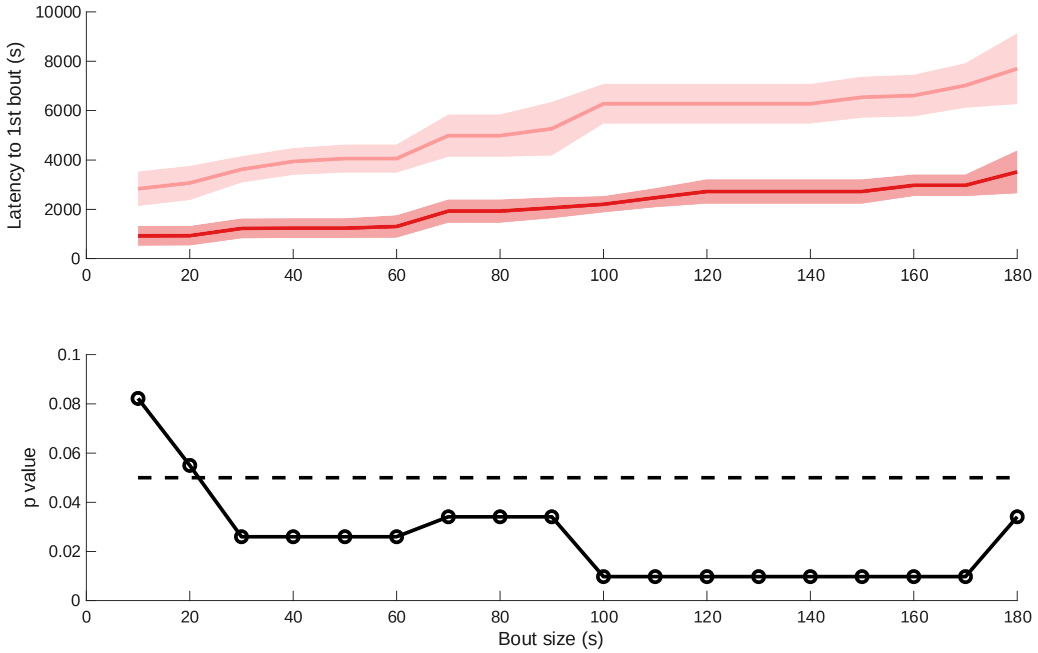


**Latency to first SWS bouts of different durations.** When compared to CONTROL (unsaturated red), LESION (saturated red) animals consistently displayed decreased values of latency to the first SWS bouts with duration ranging from 30 to 170 s. Top panel shows median S.E.M. for the different bout sizes and the bottom panel displays the p-value for each case, with the dashed line marking significance level at 0.05 (Mann Whitney U Test; Benjamin Hochberg correction for multiple comparisons).

**Fig. S2.**

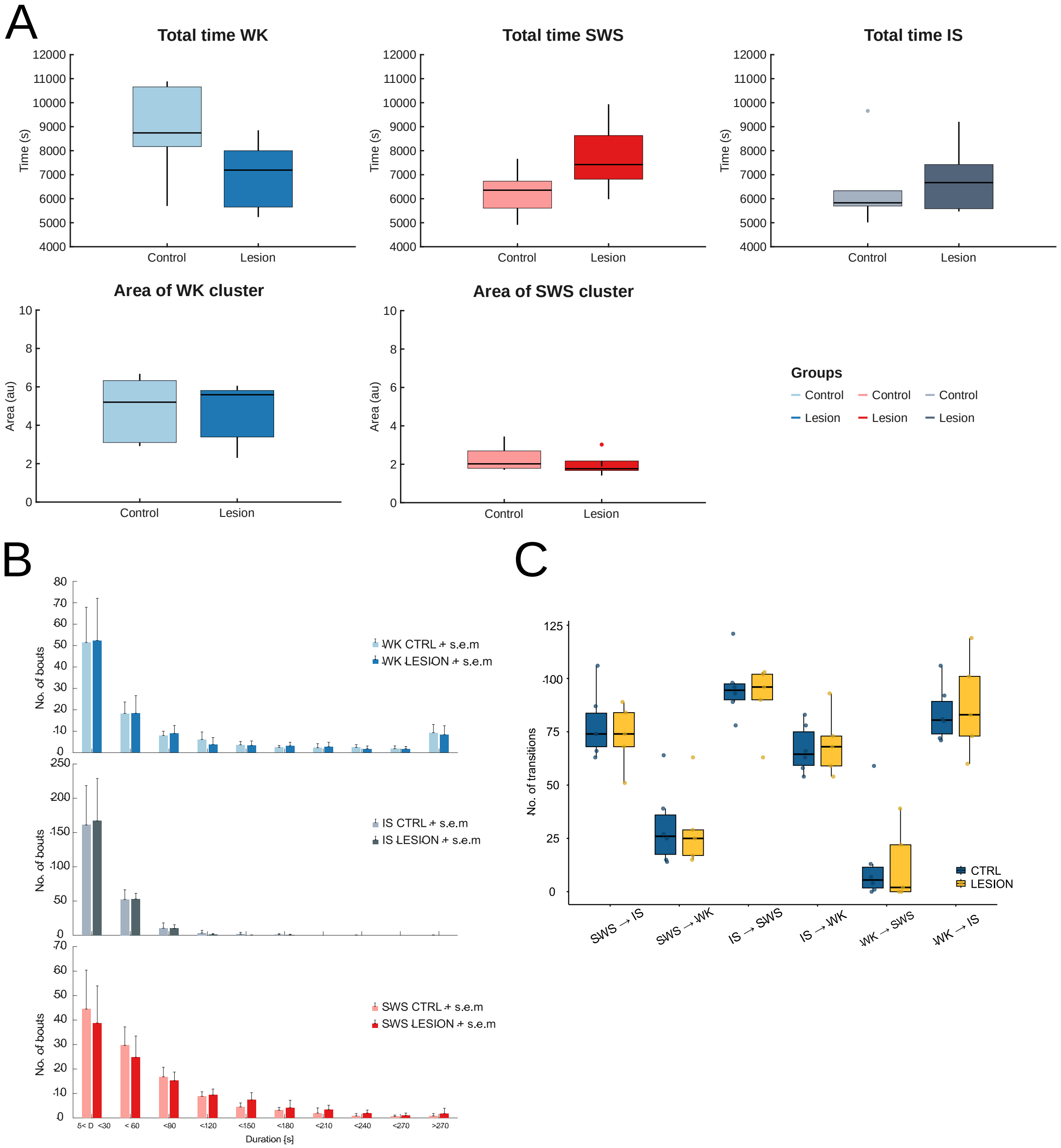


**Hypnogram statistics.** (A) Total time spent in each state (WK and SWS) as well as in the transition (IS) did not show any significant differences between CONTROL and LESION animals, even though there seemed to be a trend to a decreased duration of WK and increased duration of SWS in LESION (top row). The area of the clusters of each stage showed no difference between conditions (bottom row). As a transition state between stages and not properly a cluster, area of IS was not computed. (B) Sleep fragmentation, measured as the frequency of occurrence of bouts of different sizes, was not different between groups, in any of the states (WK, IS, and SWS; top, middle, and bottom rows respectively), for any of the computed durations. (C) Transitions between all and each possible pairs of stages were also not different across groups.

**Fig. S3.**

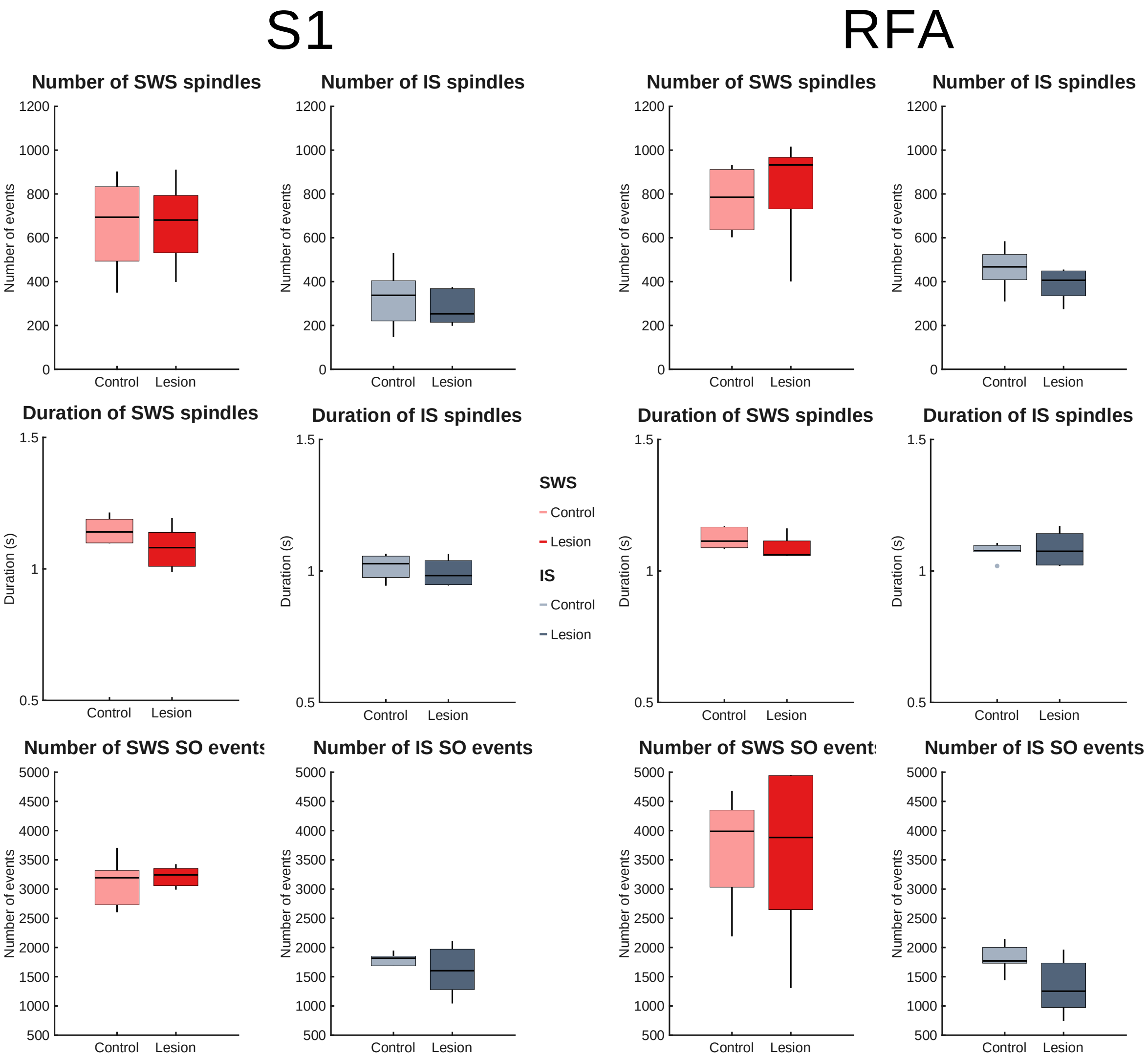


**Spindle and SO events occurrence.** This plot shows the number of spindles (top row), their duration (middle row), and the number of SO events (bottom row) during SWS (red) and IS (grey). The two leftmost columns relate to the S1 area and the two rightmost columns are RFA (two rightmost columns). Although a trend of a decreased number of spindles and duration in LESION (saturated colours) when compared to CONTROL animals (unsaturated colours) was noticeable in many instances, there were no significant differences detectable.

**Fig. S4.**

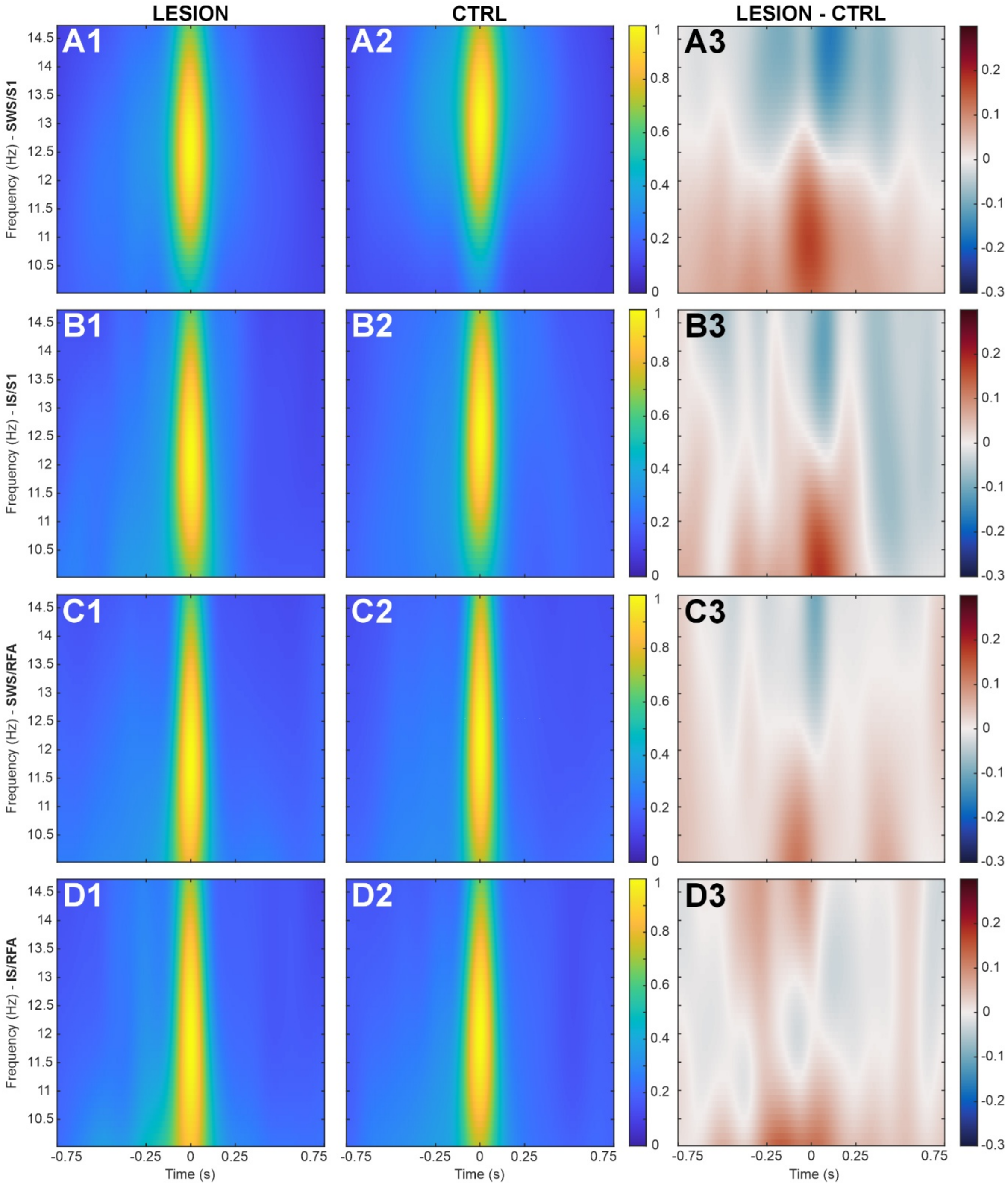


**Spindle spectral content reflects stage-specific effects of the ischemic lesion**. **Columns 1 (A1, B1, C1, D1) and 2 (A2, B2, C2, D2)**: average Continuous Wavelet Transform (CWT) spectrograms of spindles activity in the Local Field Potentials (LFPs). Columns 1 and 2 portray the group-average spectrogram (for CTRL and LESION group, respectively) of the spindle events occurred in the longest bout of a given sleep stage, either Slow Wave Sleep (SWS) (Rows A and C) and Intermediate Sleep (IS) (Rows B and D). Spectrogram power scale has been normalized with respect to the maximum power value and displayed through a colormap. Results are shown for both the implantation areas, primary somatosensory cortex (S1;

Rows 1 and 2) and Rostral Forelimb Area (RFA) (Rows 3 and 4). All the spectrograms are shown with bicubic spline interpolation for visualization only. **Columns 3 (A3, B3, C3, D3)**: Differential spectrograms obtained as the subtraction LESION – CTRL for each of the sleep stage/area (i.e., Column 1 – Column 2). Differences between groups are shown with their polarity via colormap, with red and blue representing positive and negative differences respectively. As for spectrograms, interpolation was used for visualization only.
